# HIV-1 single transcription start site mutants display complementary replication functions that are restored by reversion

**DOI:** 10.1101/2024.12.04.626847

**Authors:** K. GC, S. Lesko, A. Emery, C. Burnett, K. Gopal, S. Clark, R. Swanstrom, N.M. Sherer, A. Telesnitsky, S. Kharytonchyk

**Affiliations:** Department of Microbiology and Immunology, University of Michigan Medical School, Ann Arbor, Michigan, USA; McArdle Laboratory for Cancer Research, Institute for Molecular Virology, & Carbone Cancer Center, University of Wisconsin, Madison, WI, USA; UNC Lineberger Comprehensive Cancer Center, University of North Carolina at Chapel Hill, Chapel Hill, North Carolina, USA; Department of Biochemistry and Biophysics, University of North Carolina at Chapel Hill, Chapel Hill, North Carolina, USA; UNC Center for AIDS Research, University of North Carolina at Chapel Hill, Chapel Hill, North Carolina, USA; Cellular and Molecular Biology Program, University of Michigan Medical School, Ann Arbor, Michigan, USA

## Abstract

HIV-1 transcription initiates at two positions, generating RNAs with either ^cap^1G or ^cap^3G 5′ ends. The replication fates of these RNAs differ, with viral particles encapsidating almost exclusively ^cap^1G RNAs and ^cap^3G RNAs retained in cells where they are enriched on polysomes and among spliced viral RNAs. Here, we studied replication properties of virus promoter mutants that produced only one RNA 5′ isoform or the other: separately, in combination, and during spreading infection. Results showed that either single start RNA could serve as both mRNA and genomic RNA when present as the only form in cells, although ^cap^3G RNA was more efficiently translated and spliced while ^cap^1G RNA was packaged into nascent virions slightly better than RNAs from the parental virus. When co-expressed from separate vectors, ^cap^1G RNA was preferentially packaged into virions. During spreading infection ^cap^1G-only virus displayed only minor defects but ^cap^3G-only virus showed severe replication delays in both the highly permissive MT-4 cell line and in primary human CD4+ T cells. Passage of ^cap^3G-only virus yielded revertants that replicated as well as the twinned (^cap^1G+ ^cap^3G) transcription start site parent. These revertants displayed restored packaging and splicing levels and had regained multiple transcription start site use.

**Importance:** HIV-1 generates two RNAs during its replication that differ by only two nucleotides in length. Despite this very minor difference, the RNAs perform different and complementary replication functions. When mutants that expressed only one RNA were forced to revert, they regained functions associated with the second RNA.

## Introduction

The generation of HIV-1 RNAs requires recruitment of host RNA polymerase II to a single transcriptional promoter on integrated DNA. After transcription, a subset of HIV-1 RNAs is exported from the nucleus without first being spliced while other RNAs undergo alternative splicing to produce multiple viral mRNA species. Unspliced viral RNA plays two roles essential for viral replication: serving as mRNA for viral Gag and Gag-Pol polyproteins or becoming encapsidated into nascent virions as viral genomic RNA (gRNA).

Recently it was shown that the HIV-1 possesses a twinned promoter, with transcription initiating at two distinct transcription start sites (TSS) separated by two nucleotides [1, 2]. As a result, two major viral precursor mRNAs are formed, ^cap^1G RNA and ^cap^3G RNA, that differ in length by two nucleotides at their 5′ ends. Surprisingly, this heterogeneous transcription initiation is a major determinant of function for these two viral RNAs. Specifically, essentially all the viral RNA packaged into virions has ^cap^1G ends, whereas ^cap^3G RNAs are enriched among viral mRNAs associated with polysomes and spliced viral mRNAs [1–3].

Despite differing by only two nucleotides, the 5′ leaders of the two primary transcripts can adopt drastically different structures [4, 5] (Fig.1A). The ^cap^3G conformer predominantly folds such that the dimerization initiation site (DIS) RNA palindrome is sequestered in a double-stranded region, whereas both the major 5′ splice site and 5′ cap structure are exposed. The accessibility of these elements is reversed in the conformer most readily adopted by HIV-1 ^cap^1G RNA, with the DIS loop exposed and the splice donor signal and 5′ cap sequestered by intramolecular interactions. Accessibility of the DIS loop is required for gRNA dimerization and packaging [6, 7]. Thus, the folding propensities of the two RNA isomers suggest that inefficient ^cap^3G RNA packaging reflects the sequestration of its DIS, while splicing and translation are enabled by the accessibility of the splicing signal and 5’-cap [4]. Consistent with this model, RNAs with ^cap^3G leaders fail to form dimers but efficiently bind the translation initiation factor eIF4E *in vitro*, whereas ^cap^1G RNAs dimerize efficiently but bind eIF4E inefficiently [4, 5].

**Fig. 1.**
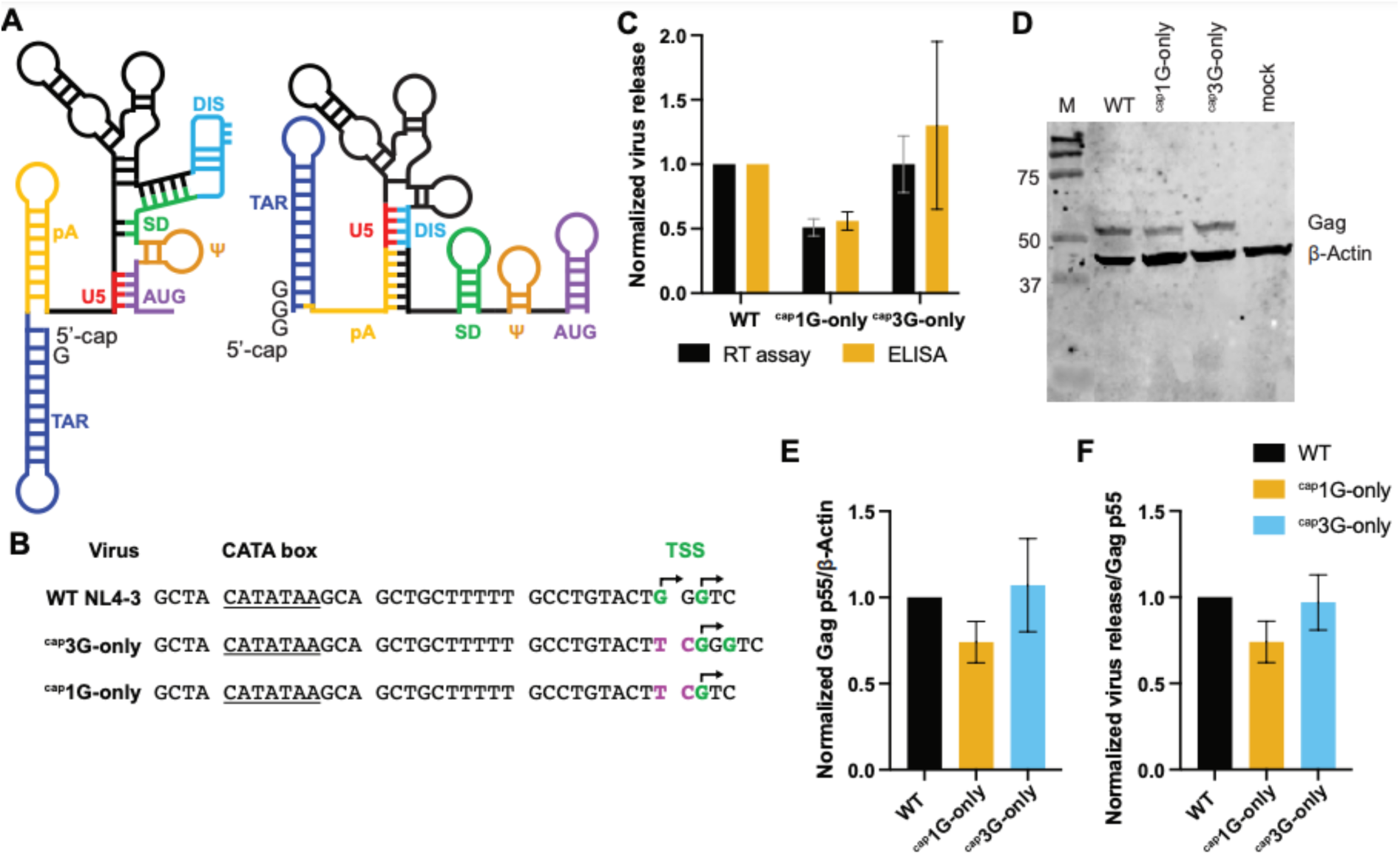
Both HIV-1 RNA 5′ end isoforms can serve as mRNA. (A) Predominant secondary structures ^cap^1G (right) and ^cap^3G (left) HIV-1 5′-leader RNAs. Sequence motifs (indicated by U5, DIS, SD, etc. [5]) are shown in separate colors that are the same in both conformers, to indicate that identical sequences form alternate structure elements. (B) Core promoter elements, including CATA box and TSS, in parental NL4-3 strain HIV-1 (WT) and in ^cap^3G- and ^cap^1G-only mutant promoters. WT start sites are indicated in green, insertions/substitutions in the single TSS mutants are indicated in purple, mapped TSS [8] are indicated with arrowheads (C) Virus release levels from transfected 293T cells quantified by RT activity or p24 ELISA, normalized to WT levels set to 1; (D) Gag examined by western blot analysis (E) Calculated p55 Gag/b-actin ratios (F) Virus release per unit Gag, based on data in panels C and E. Data in panels C, E, and F were from three independent experimental replicates.

Recent studies have mapped the HIV-1 core promoter determinants of heterogeneous transcription initiation [8, 9]. These findings showed that sequences adjacent to the TSS and the distance between the CATA-box element and TSS play crucial roles in twinned transcription initiation. This led to the identification of HIV-1 promoter mutants with focused TSSs that initiate transcription from a single position [8]. Viruses generated by these mutants, which produce only ^cap^1G RNAs or only ^cap^3G RNAs, show differing levels of replication deficiency in CEM-SS cells. Specifically, whereas ^cap^3G-only virus displays severe defects when compared to the parental virus, ^cap^1G-only virus replicates only slightly less well than its wild type (WT) twinned TSS parent [8]. Note that throughout the current report, ‘WT’ is used as shorthand for the parental NL4-3 strain of HIV-1, with its twinned TSS promoter that produces both ^cap^1G and ^cap^3G RNAs, and the virions it produces.

In the current study, the nature of replication defects of the single TSS mutant viruses were examined. We determined that ^cap^1G-only and ^cap^3G-only mutant viruses differ in RNA packaging, splicing, and translation efficiency. We compared the replication properties of wild type and mutants in highly permissive MT-4 cells and in human primary CD4+ T cells and also selected for revertants. Several revertants of the highly defective ^cap^3G-only virus were isolated that displayed restored replication efficiency. Analysis of these revertants revealed that each had recovered the ability to generate multiple RNA 5′ isoforms, displayed improved packaging, and had restored splicing levels.

## Results

### Both HIV-1 RNA 5′ isoforms can serve as mRNAs

Single TSS promoter mutants (Fig.1B) were introduced into a replication defective vector that included the HIV-1 RNA leader, the *gag, gag-pol, tat* and *rev* genes, and a puromycin resistance expression cassette in place of portions of *env* [5]. Both ^cap^1G-only and ^cap^3G-only derivatives produced viral particles upon transient transfection, but ^cap^3G-only virus yields were ∼2-fold higher than the ^cap^1G-only vector (Fig.1C). Intracellular Gag levels were compared by western blot analysis (Fig. 1D). After normalizing to β-actin, the data revealed that cells with the ^cap^3G-only vector contained ∼1.5-fold more intracellular Gag than ^cap^1G-only, and that ^cap^3G-only Gag levels were similar to those of WT (Fig.1E). When normalized to intracellular Gag, virion release for WT and ^cap^3G-only were indistinguishable, with a possible minor but not significant decrease in virion release by ^cap^1G-only (Fig.1F).

In summary, both HIV-1 RNA isoforms can serve as mRNAs when they are the only RNA form in cells. However, consistent with previous findings of an enrichment of ^cap^3G RNA on polysomes [1], more Gag protein was produced and more virions were released by the ^cap^3G-only mutant compared to the ^cap^1G-only mutant.

### Both RNA isoforms can be packaged and serve as genomic RNAs, albeit with diCering eCiciencies

Next we examined the extent to which each RNA isoform could be packaged when it was the only RNA present. Cells were transiently transfected with either the WT or the single TSS vectors described above. An RNase protection assay (RPA) was performed to compare levels of viral RNA (annealed to a probe within the *gag* gene) relative to the amount of the host 7SL RNA, which is packaged into virions in proportion to the viral Gag protein (Fig. 2A) [10]. The results revealed that ^cap^1G-only RNA was packaged slightly (∼1.2-fold) better than RNAs generated by the WT vector. In contrast, packaging of ^cap^3G-only RNA was reduced ∼1.6-fold relative to WT vector RNAs, indicating that ^cap^3G RNAs were packaged ∼2-fold less efficiently than ^cap^1G RNAs (Fig. 2B).

**Fig. 2.**
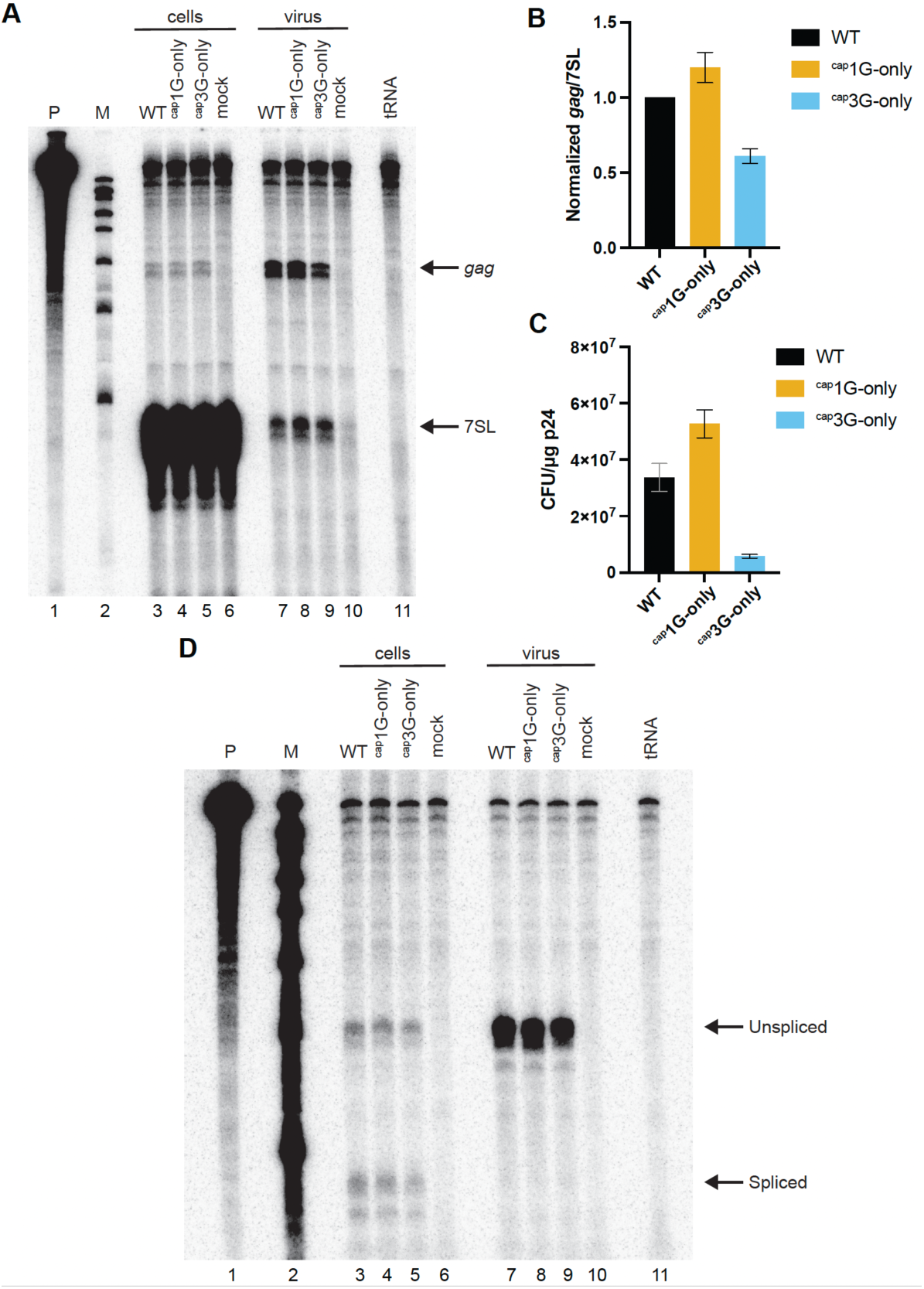
Both 5′ end isoforms can be packaged and serve as gRNA. (A) RNAse protection assay (RPA) of viral RNA in transfected 293T cells and virions. Probe fragments protected by HIV-1 vector RNAs *gag* and the host normalization standard 7SL RNA (7SL) are indicated. Cell samples are at the left and virion RNAs are on the right. Lane designations indicate transfected vectors; Mock: mock-transfected cells; tRNA: yeast tRNA control; Ladder: molecular weights marker; Probe: undigested chimeric *gag*-7SL riboprobe. (B) RNA packaging ebiciencies. Using RPA data quantified by phosphorimager analysis, RNA levels were first normalized to 7SL levels, then virion values were divided by cell RNA levels, with the WT sample assigned a value of 1. (C) Puromycin resistant colony forming titers. Titers were determined for WT NL4-3 GPP vector and single TSS NL4-3 GPP vectors pseudo-typed with VSVG envelope (see Materials and Methods). The Y axis indicates cfu titers per 1μg of HIV-1 p24 as determined by RT-activity levels on infections using virus from three independent transfections. (D) Spliced viral RNA production and packaging in the cells transfected with NL4-3 GPP derivative vectors determined by the RPA. Riboprobe HIV unspliced/spliced (see Materials and Methods) was used in this experiment. RNA samples extracted from cells are at the left and those from virus-containing media are on the right. Migration positions of protected fragments are indicated on the right.

Because HIV-1 virions ordinarily package ^cap^1G RNAs, this is the only 5’ isoform delivered to newly infected cells. To test if early replication steps were as efficient for ^cap^3G RNAs as for ^cap^1G RNAs, encapsidated WT, ^cap^1G-only, and ^cap^3G-only vectors were tested in a single cycle infectivity assay. When normalized by the levels of reverse transcriptase activity (RT) in the medium, viral particles generated by the ^cap^1G-only vector showed a ∼1.5-fold higher puromycin resistant colony forming unit titer than those from the WT vector (Fig. 2C) – a value similar to the ∼1.2-fold higher level of ^cap^1G-only vector RNA packaging observed above (Fig. 2B). However, virions from the ^cap^3G-only vector showed an approximately 6-fold lower titer than virions produced by the WT vector. This result indicates that ^cap^3G-only vectors have replication defects in addition to their modest packaging defects. This early defect is consistent with a defect in viral DNA synthesis, and it has recently been shown that ^cap^3G RNAs serve less efficiently as reverse transcription templates than ^cap^1G RNAs, both *in vitro* [2] and during viral replication [10]. Also of note, although WT virus predominantly packages ^cap^1G RNA, about 15% of 293T cell-produced WT virions contain gRNAs with alternate 5′ end sequences [9, 11]. It is conceivable that the minor enhancement in ^cap^1G-only virus titer relative to WT is due to an absence of alternative 5′ isoform packaging by this mutant relative to the low level of non-^cap^1G RNA packaged by the WT.

HIV-1 RNA packaging is notoriously promiscuous, in that RNA packaging element mutants are well-packaged in the absence of WT competition [12–16]. Furthermore, whereas most viral RNA in HIV-1 particles is unspliced and full-length, a small amount of packaged spliced RNA can be detected, and spliced RNA packaging increases for mutants with RNA dimerization and encapsidation defects [6, 17, 18]. Thus, to address the possibility that some of the observed defects in ^cap^3G RNA packaging might reflect enhanced spliced viral RNA packaging, cell and virion RNAs were compared by RPA using a probe that spans the major 5′ splice site, D1 (Fig. 2D). The results indicated that spliced and unspliced viral RNAs were readily detected in cells for each single TSS mutant and the WT vector, and whereas modest differences in ^cap^3G and ^cap^1G RNA splicing levels have been reported [3], these differences were not apparent by the less quantitative RPA approaches used here (Fig.2D, cells). In contrast, only unspliced RNAs were detected in virion RNA samples (Fig. 2D, virus). Thus, the observed diminution of packaging for ^cap^3G-only was not a result of excessive spliced RNA packaging, and packaging specificity for full-length gRNA was retained by the ^cap^3G-only mutant.

### Single virion analysis confirmed high-level ^cap^1G RNA packaging

Previous imaging work has shown that viral RNA is detectable in >90% of HIV-1 virions produced by transfected cells, leaving open the possibility that a minor fraction of virions does not contain gRNA [6]. Thus, a single viral particle fluorescent microscopy assay was performed to address if the apparent elevated level of ^cap^1G gRNA packaging per unit virion protein observed above corresponded to a higher proportion of gRNA-containing viral particles for the ^cap^1G-only vector than for WT.

This analysis was achieved using a series of self-labeling Gag-YFP/MS2-mCherry reporter viruses. Using these, particles were visualized by YFP and viral RNA was detected by the presence of an MS2-mCherry fusion protein, which bound to MS2 binding sites on the viral RNA (Fig. 3A). Collecting YFP and mCherry channels and finding colocalization of YFP and mCherry signals in a cell-free punctum was indicative of a virion containing gRNA. As a control, viral particles that had a deletion in NC, a viral protein needed in packaging (ΔNC), showed no YFP colocalization with the mCherry signal.

**Fig. 3.**
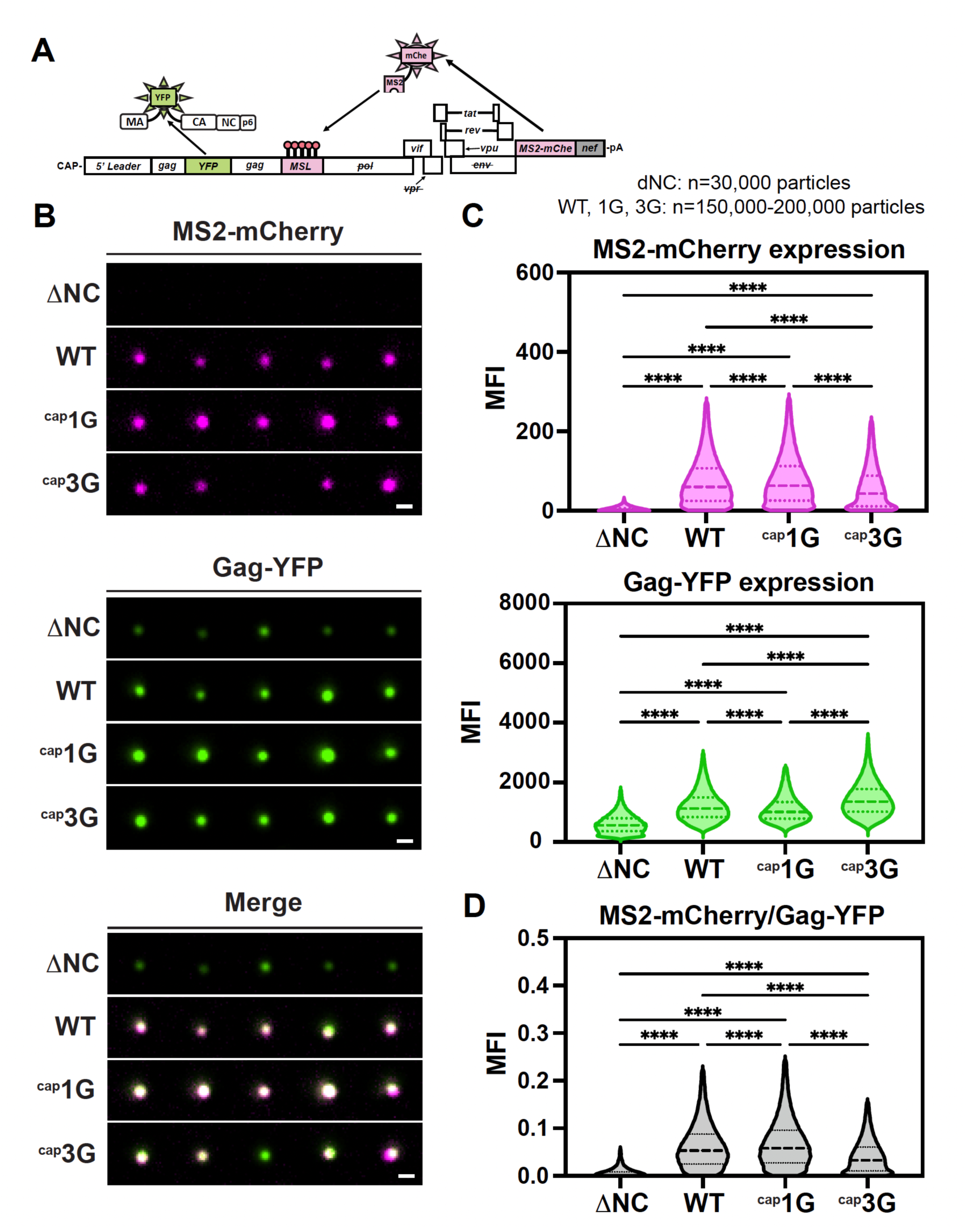
Single-virion analysis shows virions from ^cap^1G-only virus display a higher packaging eIiciency than ^cap^3G and WT. (A) Schematic representation of the two-color self-tagging reporter virus (pNL4-3 Gag-mVenus/ 24xMSL/MS2-mCherry). (B) Representative images of single fluorescent virions harvested from transfected HEK 293T cells. Scale bar = 0.5μm. DNC, reporter virus with WT promoter and deletion of the NC domain of the Gag; WT, virus with WT promoter, ^cap^1G, and ^cap^3G, reporter viruses with corresponding single TSS mutations. (C) Quantification of single virions for DNC, WT, ^cap^1G, and ^cap^3G viruses showing ratio of virions with MS2-mCherry and Gag-YFP mean fluorescent intensities (MFIs), as a ratio of WT. (D) MS2-mCherry signal per Gag-YFP MFI for DNC, WT, ^cap^1G, and ^cap^3G virions. For all violin plots (C and D), dashed lines indicate median and dotted lines indicate 25th and 75th quartiles. ****P<0.0001.

Virions produced by transfection were purified, plated on glass coverslips, and imaged on a widefield microscope (Fig. 3B). When ratios of mCherry to YFP per punctum were determined, the results showed a slight (∼1.1-fold) increase in MS2-mCherry labeled RNA per punctum for ^cap^1G-only relative to WT virus (Fig.3C, D, Table S1). Consistent with the RPA data above, an ∼1.7-fold increase in signal colocalization relative to ^cap^3G-only virus was observed (Fig. 3C, D, Table S1). Taken together, mCherry to YFP levels per punctum were reduced ∼1.5 and ∼1.7-fold in ^cap^3G-only viral particles relative to WT and ^cap^1G-only viruses, respectively (Fig.3D, Table S1). The 1.1-fold increase in Gag and viral RNA signal co-localization for the ^cap^1G-only virus was close to the 1.2-fold increase in viral RNA packaging observed by RPA, providing further evidence that a larger proportion of ^cap^1G-only viral particles contain viral RNA than do WT virions. Similarly, the 1.5-fold decrease in YFP and mCherry co-localization relative to WT matched the 1.6-fold decrease in ^cap^3G-only virus RNA packaging measured by RPA above.

### ^cap^1G RNAs readily out-compete ^cap^3G RNAs for packaging

The work above examined 5′ isoform properties when each was the only HIV-1 RNA present in cells. However, because both RNAs are present during natural HV-1 infection, experiments were also performed where the two single TSS mutants were co-expressed. A packaging-defective Ψ-helper that provided all HIV-1 proteins in *trans* was used to mobilize TSS mutant vectors in which all HIV-1 coding regions were deleted (Minimal vectors) [5]. Viral particles were harvested from cells co-transfected with Ψ-helper plus pairs of Minimal vectors, and cell and virion RNA was assayed by RPA (Fig. 4A, B). In each Minimal vector co-transfection, one of the two (Minimal Δ) contained a deletion in sequences that do not contribute to packaging specificity. As a result, Minimal and Minimal Δ vector RNAs protected different-sized riboprobe fragments that allowed separate identification of the co-expressed vectors by RPA. Analysis of the RNAs in viral particles produced by co-transfected Minimal plus Minimal Δ vector pairs revealed that the presence of ^cap^1G RNA effectively prevented ^cap^3G RNA encapsidation, regardless of whether ^cap^1G was expressed by a Minimal or a Minimal Δ vector (Fig. 4A, B). Consistent with previous reports, Ψ-helper RNA was observed in virions from cells transfected with Ψ-helper alone (Fig. 4A lane 9) but all Minimal vectors efficiently outcompeted the Ψ-RNA for packaging [13–17].

**Fig. 4.**
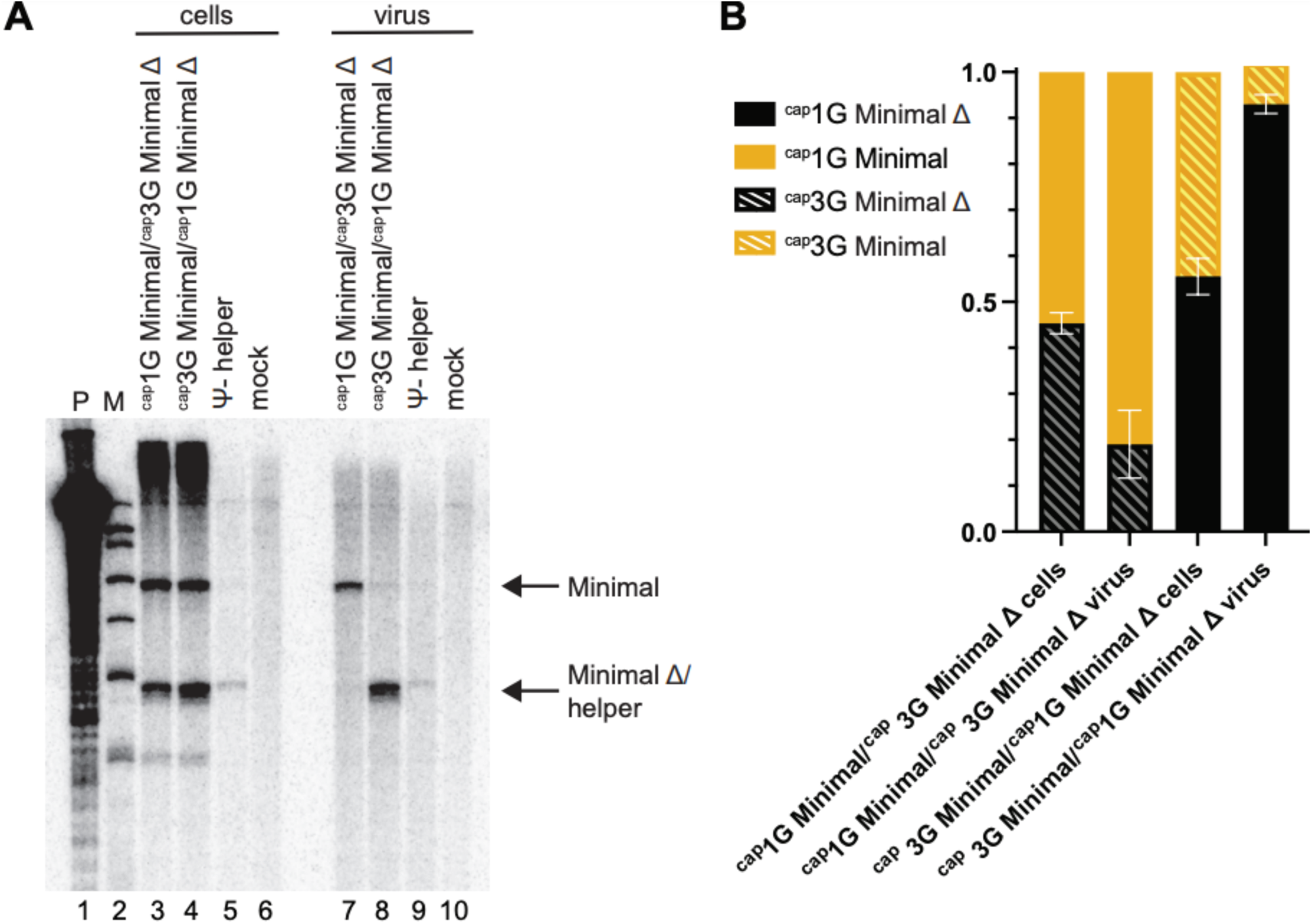
^cap^1G-only RNAs outcompete ^cap^3G-only RNAs for packaging. (A) Packaging ebiciency in competitive conditions. RPA of cell and virus samples resulting from co-expression of Y-helper with both ^cap^1G-only and ^cap^3Gonly vectors. Protected probes fragments are indicated on the right. Lane designations indicate transfected vectors. P: undigested riboprobe; M: size markers; mock: mock-transfected cells. (B) Proportions of ^cap^1G and ^cap^3G RNAs in cells and virions, as determined by RPA using RNA samples from two independent experiments.

### Peak virus levels achieved during spreading infection by ^cap^3G-only virus were lower than those of ^cap^1G-only in MT-4 cells and in primary cells

Replication studies were performed using the infectious NL4-3 clone containing ^cap^1G-only and ^cap^3G-only mutations. Previous work with these mutants in CEM-ss cells showed that ^cap^1G-only virus replication was only minimally delayed relative to WT, and that peak levels of replication were observed 3-4 weeks post-infection. In contrast, ^cap^3G-only virus remained at low levels for the duration of these previous experiments [8].

Here, we used the highly permissive MT-4 cell line [19, 20] as well as stimulated primary CD4+ T cells to study replication kinetics and to select for revertants. After infection, culture media were sampled every 2-3 days to monitor viral particle production. At the same time points, infected cell samples were harvested for proviral DNA analysis (Fig. 5A). As previously observed using CEM-ss cells, replication kinetics of ^cap^1G-only virus were similar to but slightly slower than WT in MT-4 cells. In contrast to the previous studies, ^cap^3G-only virus did not remain at a low level but instead expanded through the culture, albeit slightly slower than and reaching a peak 2-5 days later than WT or ^cap^1G-only (Fig.5A). Similar trends were observed in primary cells, with ^cap^3G-only virus replicating slower than WT or ^cap^1G-only virus, and with ^cap^1G-only replication kinetics very similar to those of WT NL4-3 (Fig. 5B). Thus, consistent with the packaging and early replication stage defects observed above, the ^cap^3G-only virus showed reduced replication capacity when tested in a spreading viral infection.

**Fig. 5.**
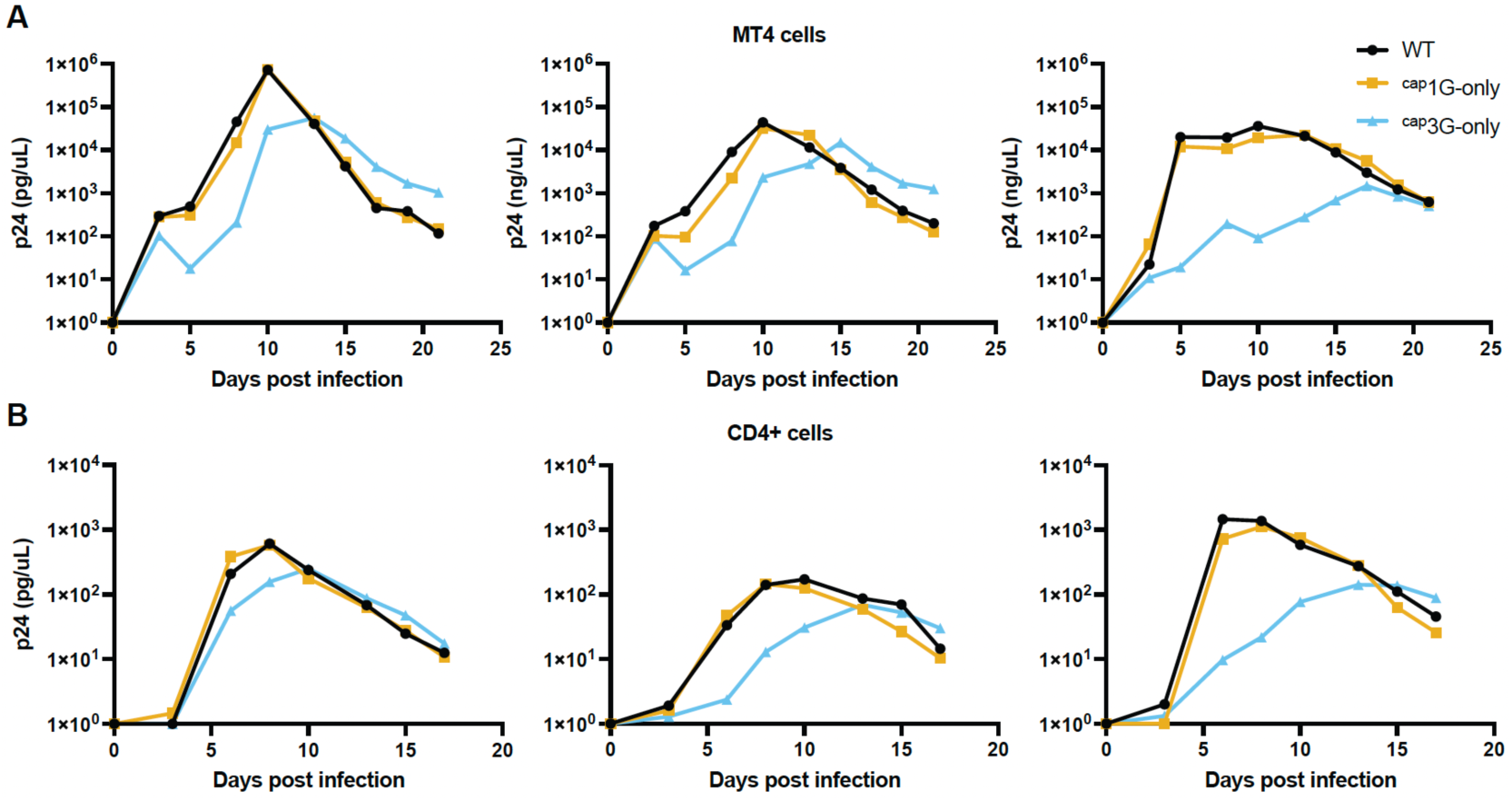
Replication kinetics of the single TSS NL4-3 infectious viruses. Replication kinetics of the NL4-3 derivatives in MT4 cells (panel A) and in the primary blood CD4+ cells (B) as monitored by quantifying media RT levels and normalizing to p24. Each graph represents one independent experiment.

### Fitness of WT and a ^cap^1G-only revertant diCered little from parental ^cap^1G-only virus

Virus evolution was tracked by sequencing cell-associated viral DNA extracted at various time points post-infection. The data revealed that a TSS region mutation (TCG to TGG, hereafter called ^cap^1G-R1) (Fig. 6A) emerged by day 3 and co-replicated with the original ^cap^1G-only virus throughout a 3-week infection (Fig. 6A). Upon diluting these cultures into fresh MT-4 cells, the proportion of ^cap^1G-R1 gradually increased, but the parental ^cap^1G-only mutant persisted throughout the 60 day-long experiment and no additional TSS revertants were observed. The same revertant emerged during passage of ^cap^1G-only virus in primary CD4+ T cells (Fig. 6B).

**Fig.6.**
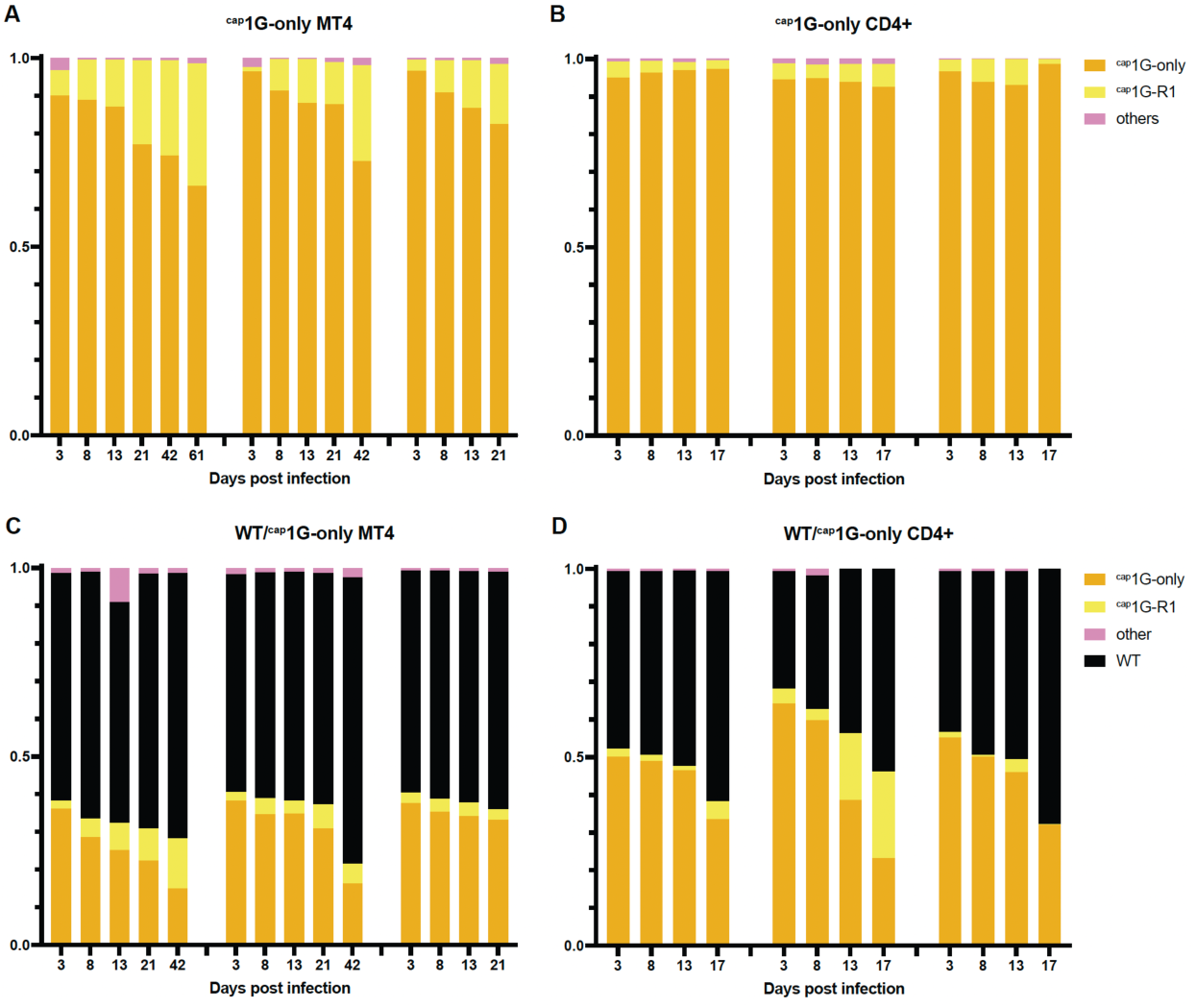
^cap^1G-only virus fitness and revertant selection. Proportions of TSS variants in ^cap^1G-only virus infected MT4 (A) or primary CD4+ cells (B) at indicated timepoints, as observed by high throughput sequencing. (C) and (D) Changes in TSS variant proportions in MT4 (C) or primary CD4+ blood cells (D) co-infected with WT NL4-3 plus^cap^1G-only virus over time, as observed by high throughput sequencing. Each graph represents one independent experiment.

To further study ^cap^1G-only virus fitness, cells were coinfected with WT NL4-3 and infectious ^cap^1G-only virus. High throughput sequencing revealed the emergence of ^cap^1G-R1 and a slow decrease in the proportion of ^cap^1G-only virus over time, but WT did not completely out-compete the ^cap^1G-only mutant or its revertant (Fig.6C, D). Together these results confirmed that ^cap^1G-only virus is only slightly less fit than WT in the cell types studied here and showed that its predominant revertant gained at most a minor amount of fitness.

### Revertants dominated cultures during spreading infection with ^cap^3G-only viruses

^cap^3G-only MT-4 cell cultures were rapidly dominated by revertants (Fig.7). High throughput sequencing revealed that by day 3, the same TSS region revertant (^cap^3G-R1; TCGGG to TGGGG) (Fig.7A, C) was detectable in all three independent infections. The proportion of this revertant increased over time while the original ^cap^3G-only variant gradually decreased (Fig.7A). Additional ^cap^3G-only revertants emerged later, including TCGGA (^cap^3G-R2) and a one base deletion revertant, TCGG (^cap^3G-R3) (Fig.7A, C).

**Fig. 7.**
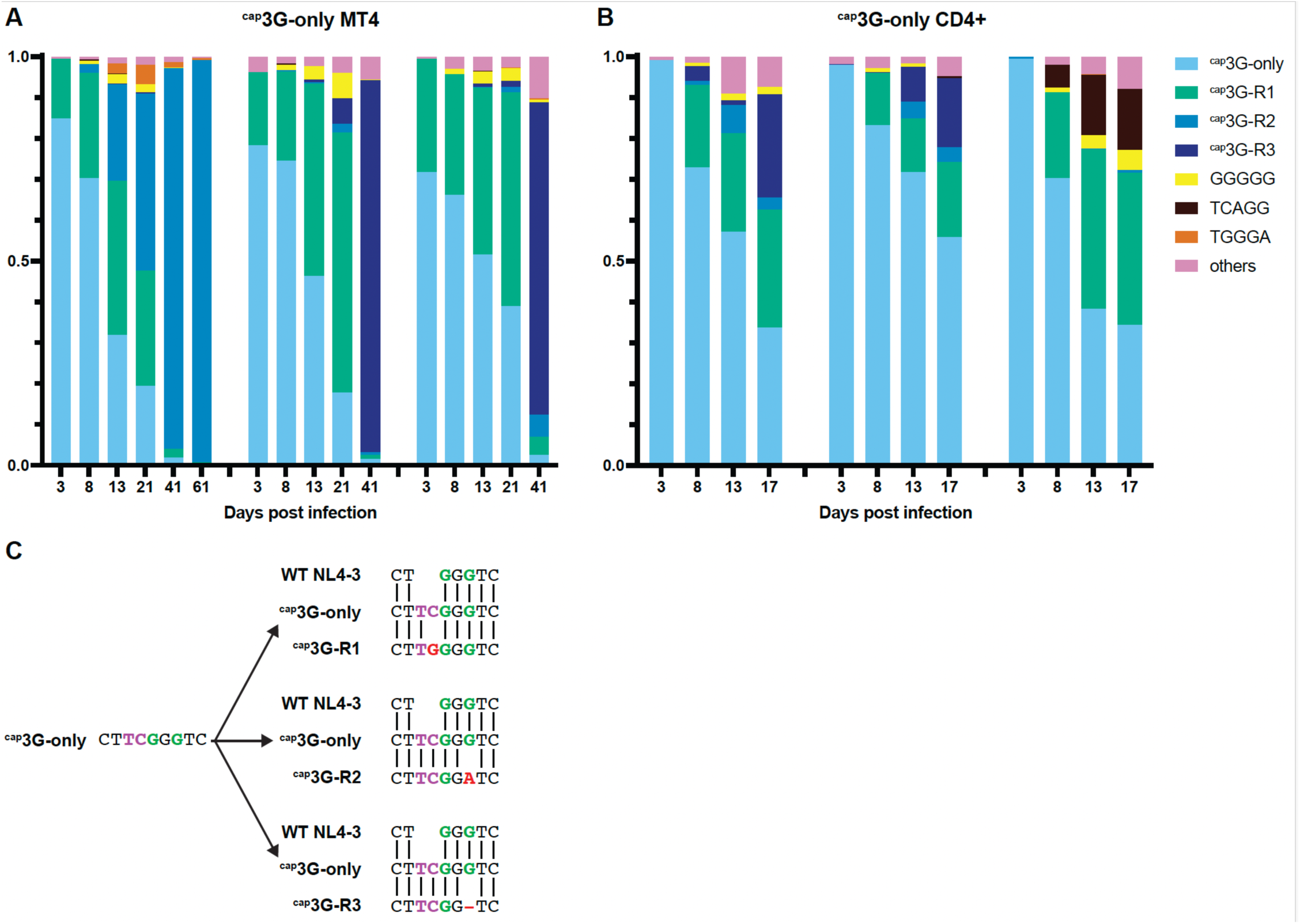
Selection of ^cap^3G-only virus revertants. Proportions of TSS variants in ^cap^3G-only virus infected MT4 (A) and primary CD4+ cells (B) over time, as observed by high throughput sequencing. Each graph represents one independent experiment. (C) Alignment of the TSS sequences of the 3 most prominent ^cap^3G-only virus revertants with the ancestral single TSS mutant and WT NL4-3.

Rarer variants were observed episodically, with the most prominent (GGGGG and TGGGA) reaching about 5% of the population at intermediate time points but then largely disappearing at later time points (Fig.7A). Some late timepoint subclones displayed additional mutations outside the TSS region, but these did not become fixed in the populations and most late timepoint proviruses carried single mutations in their TSS regions only (Supplementary data). No changes in *gag* were observed.

All three major ^cap^3G-only revertants that emerged in MT-4 cells were also observed during infection of primary cells (Fig.7B). Population dynamics appeared less consistent in primary cells than in MT-4 cells, including the emergence of several additional variants (including one with TSS sequence TCAGG, which reached about 15% of the third replicate’s population) that were not observed in MT-4 cells (Fig.7B).

### ^cap^3G-only revertants generated multiple RNA isoforms, showed improved fitness, and displayed restored levels of packaging and splicing

The rapid dominance of revertants in ^cap^3G-only cultures suggested that these variants had acquired fitness advantages. To assess this, the revertants ^cap^3G-R1, ^cap^3G-R2 and ^cap^3G-R3 were built into lentivirus vectors and infectious molecular clones to test replication properties and investigate if the changes restored functions that had been rendered defective by ^cap^3G-only mutations. One prominent difference between WT and ^cap^3G-only virus is that replication of the former generates two RNA 5’ isoforms and the latter only one. Thus, the 5′ ends of virion RNA were mapped at single base resolution (Fig.8A).

**Fig. 8.**
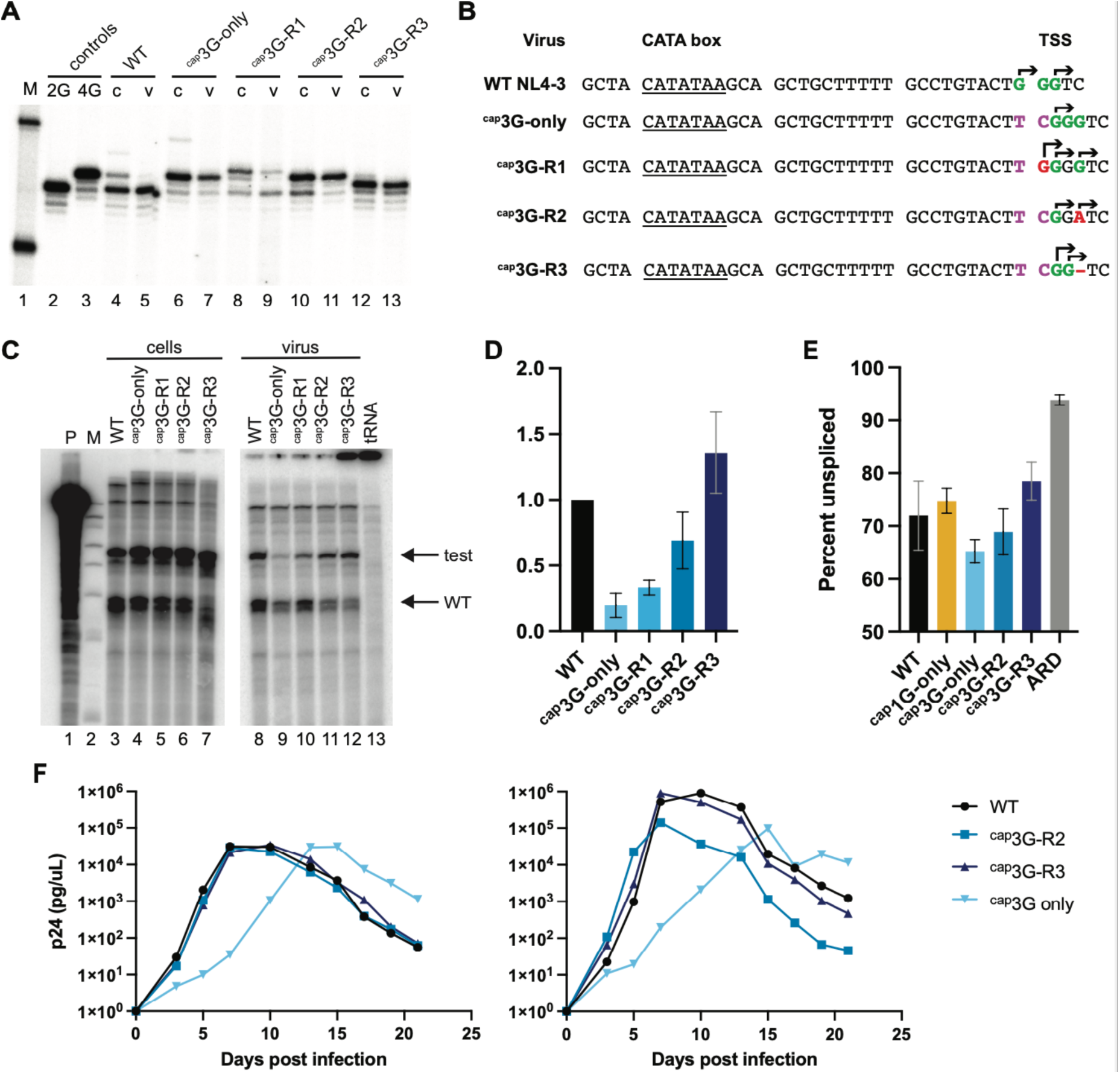
^cap^3G-only revertants restore RNA functions and replication capacity. (A) Single base resolution assay RNA 5′ ends produced by ^cap^3G-only revertants. M: size markers; controls: 2G and 4G controls, which migrate at the positions of ^cap^1G and ^cap^3G products, respectively; RNA samples from cells (lanes indicated “c”) and virus (v). (B) Core promoter of NL4-3, ^cap^3G-only mutant and revertants. CATA box underlined. The two major WT TSS are in green. Observed TSSs indicated with arrows. TC insertion responsible for the ^cap^3G-only phenotype shown in purple. Mutations in the revertants shown in red. (C) Packaging of revertants in competition with Ψ+ helper. Cell RPA samples at left and virus at right. Lane designations indicate vectors co-transfected with Y+ helper; P: undigested probe; M: size markers; tRNA: tRNA only control. (D) Packaging ebiciency of the revertants. Calculated by dividing the ratio of vector to helper RNA in virions by the ratio in cells. Data from two independent transfection experiments. (E) Unspliced fraction of intracellular viral RNA, as assessed by high throughput sequencing (see Methods). ARD indicates cells infected in the presence of antiretroviral drugs and confirms predicted low-level unspliced background from input virions. Results from three independent infection replicates (F). Replication kinetics of infectious ^cap^3G-only revertant clones in MT4 cells. Monitored by viral medium RT activity. Each graph represents one independent experiment.

The results revealed that the ^cap^3G-R1 revertant produced three RNA 5′ end isotypes: ^cap^4G, ^cap^3G, and ^cap^1G RNAs, with ^cap^4G the most abundant RNA in cells, followed by ^cap^1G (Fig.8A, B). In ^cap^3G-R1 viral particles, the ^cap^4G/^cap^1G RNAs’ ratio shifted toward the ^cap^1G form, although ^cap^4G and ^cap^3G RNAs were also detectably packaged (Fig.8A). The production of ^cap^1G RNA by ^cap^3G-R1, which is identical in sequence to the isoform packaged by WT HIV-1, may explain the rapid spread of this revertant in cell culture. The ^cap^3G-R2 revertant produced two RNA forms in the cells, ^cap^GGA and ^cap^A RNAs, which differed in sequence but were equal in length to the ^cap^3G and ^cap^1G RNAs of WT, respectively (Fig.8A, B). Surprisingly, the ^cap^GGA RNA isotype was the predominant form in viral particles produced by this revertant. The ^cap^3G-R3 revertant produced two RNA forms, ^cap^2G and ^cap^1G RNAs, with ^cap^2G RNA being the major RNA in cells. ^cap^3G-R3 packaged both of these RNA isoforms, with their ratio shifted toward to ^cap^1G RNA in virions (Fig.8A, B).

Because alternate 5′ ends enable HIV-1 RNAs to adopt structures required for packaging, the restoration of heterogeneous TSS usage by all ^cap^3G-only revertants suggested that their RNA packaging functions might be improved. Thus, packaging for each revertant was tested in competition with WT HIV-1. As shown in Fig. 8C and D, all three revertants displayed improved packaging compared to the parental ^cap^3G-only virus. While WT vector RNAs largely outcompeted ^cap^3G-only virus RNA in packaging, all three revertant RNAs showed an increased presence in viral particles. The revertant that generated the most competitive RNA was ^cap^3G-R3, while the ^cap^3G-R1 RNAs were the least well packaged among the revertants (Fig.8D). This higher efficiency of packaging by ^cap^3G-R2 and ^cap^3G-R3 may be part of why these revertants displaced the initial revertant, ^cap^3G-R1, during prolonged passage.

Next, splicing was addressed more quantitatively than in the experiment shown in Fig.2. Previous work has shown the enrichment of ^cap^3G-5′ ends among some classes of HIV-1 spliced RNAs [3]. Here, MT-4 cells were infected with ^cap^1G- or ^cap^3G-only mutants or with the ^cap^3G-R2 and ^cap^3G-R3 revertants, and high-throughput analysis was performed on cellular RNA to study the effects of single TSS mutations and their reversion on splicing [21]. The data revealed that unspliced HIV-1 RNA was about 72% of the total viral RNA in cells infected with WT NL4-3 virus, whereas the proportion of unspliced RNA in ^cap^1G- and ^cap^3G-only viruses was about 75% and 65% correspondingly (Fig. 8E). Thus, ^cap^3G-only had about 90% as much unspliced RNA as WT and ^cap^1G-only unspliced RNA levels were about 104% those of wild type (Fig.8E). For the ^cap^3G-only revertants, splicing was reduced and became more similar to WT, with unspliced RNA levels for ^cap^3G-R2 and ^cap^3G-R3 revertants 96% and 108% those of WT respectively (Fig.8E). Among RNAs that were spliced, the distribution of 3′ splice site usage was largely similar among variants (Supplementary Fig. 2).

Overall replication fitness of these revertants was tested in infection assays. HIV-1 NL4-3-based infectious molecular clones were generated that contained TSS region sequences from ^cap^3G-R2 or ^cap^3G-R3 in both LTRs, and these were used to generate virus stocks in 293T cells. Replication kinetics of these revertants were compared to WT and ^cap^3G-only viruses (Fig. 8F). The results indicated that both ^cap^3G-R2 and ^cap^3G-R3 replicated without the delay associated with the original ^cap^3G-only mutant and exhibited replication kinetics similar to those of WT NL4-3 virus (Fig. 8F).

## Discussion

Here we compared the abilities of the two primary isoforms of HIV-1 RNA to provide specific replication functions. We confirmed that viruses with either one of the two RNA forms alone were capable of completing a replication cycle, although ^cap^3G-only virus was much less replication-competent than ^cap^1G-only virus. At least three crucial functions of full-length viral RNA were affected in the single TSS mutants: packaging, translation and splicing. Interestingly, virus encoded by the parental NL4-3, with its twinned TSS promoter, had splicing, packaging and *gag* expression phenotypes intermediate to those of ^cap^1G-only and ^cap^3G-only viruses, suggesting that the WT phenotype is specified by the presence of its mixed RNA population.

Packaging may be the replication property most reliant on a specific RNA isoform, as WT HIV-1 displays high packaging specificity for ^cap^1G RNAs. Packaging of ^cap^3G RNA was 2-fold less efficient than ^cap^1G RNA when the RNAs were expressed separately, and ^cap^3G RNA was excluded from packaging when ^cap^1G RNA was present. Surprisingly, ^cap^1G-only viruses packaged slightly more RNA per unit virion protein than WT viruses did. Previous work has shown that >90% of HIV-1 viral particles contain viral gRNAs, thus suggesting a small fraction of particles may lack gRNA [6]. Here, packaging was measured both by determining the amount of gRNA per virion and by single virion microscopic imaging. The proportion of ‘empty’ particles that lacked gRNA, as visualized by microscopy, coincided well with changes in the packaging efficiency as measured by virion RNA quantification. The results suggest that about 10% of WT HIV-1 virions ordinarily lack gRNA, and that the proportion of “empty” virions is even lower for ^cap^1G-only viral particles.

Whereas ^cap^1G RNAs were preferentially packaged, cells transfected with ^cap^3G-only virus contained more Gag polyprotein than ^cap^1G-only-expressing cells. The 1.5-fold higher levels of intracellular Gag and increased level of virus particle release observed with ^cap^3G-only virus is consistent with reports showing that ^cap^3G RNA is translated more efficiently than ^cap^1G RNA [22, 23] and that ^cap^3G RNA is enriched on polysomes [1]. The higher levels of intracellular Gag occurs even though ^cap^3G RNA undergoes a greater level of splicing (Fig. 8E) consistent with ^cap^3G RNA being directed to translation and being less available for encapsidation. Colony forming titers per unit gRNA were similar for WT and ^cap^1G-only viruses, but titer per encapsidated gRNA was about 3-fold lower for ^cap^3G-only virus. This may reflect defects described in previous work showing that ^cap^1G RNA is more efficient as a template for reverse transcription than ^cap^3G RNA, both *in vitro* and during virus replication [2, 10].

Revertants emerged rapidly during passage of both ^cap^1G- and ^cap^3G-only viruses. Only one revertant was detected during ^cap^1G-only replication and it appeared to confer at most a very minor replication advantage. However, the ^cap^3G-only revertants replicated much better than parental ^cap^3G-only virus. For both ^cap^1G- and ^cap^3G-only viruses, the first revertants that emerged contained the same one-base substitution just upstream of the TSS. This C to G substitution at the −1 position was observed in all independent infection replicates, whether with ^cap^1G- or ^cap^3G-only viruses, and in both MT-4 and primary CD4+ T cells. Similar reversion of a ^cap^1G-only virus has been described previously [9]. The early acquisition of identical substitutions during passage of both ^cap^1G- or ^cap^ 3G-only viruses suggests that the molecular mechanism involved in reversion was the same for both viruses. It has been proposed that this reversion mutation arose during reverse transcription by the insertion of a C residue opposite the N^7^-me-G 5′-cap structure, followed by mismatch extension during later reverse transcription steps [9]. Other reports support this assertion, including findings that AMV reverse transcriptase can read through the cap *in vitro* and that C to G mutations at the −1 position are frequently observed during murine leukemia virus replication [24–26]. Rapid emergence of the −1C to G mutation during replication of both the ^cap^1G- and the ^cap^3G-only viruses suggests that HIV-1 reverse transcriptase also can efficiently read through cap residues during reverse transcription.

If cap readthrough occurs readily as a part of minus strand transfer, it seems feasible that the minor GGGGG ^cap^3G-only revertant resulted from a stepwise process involving mutations during two different rounds of replication. Specifically, this mutation, which was observed in both MT-4 and primary CD4+ T cells, might have arisen via two sequential cap read-through events: the first creating ^cap^3G-R1 with TGGGG and the second resulting during subsequent rounds of replication after the (relatively rare) packaging of a ^cap^4G RNA, followed by cap readthrough and mismatch extension. In light of this interpretation, it is interesting that the ubiquitous ^cap^1G-only virus revertant, ^cap^1G-R1 with its TGG TSS, was not observed to evolve into a WT GGG sequence through cap-copying in a subsequent replication cycle. However, replication differences between ^cap^1G-only, ^cap^1G-R1, and WT viruses are negligible, the TGG revertant principally packages ^cap^1G RNA [9], and ^cap^1G RNAs are better reverse transcription templates [10]. Together, these observations may explain why restoration of the WT TSS was not observed here or previously [9].

The rapid emergence and expansion of ^cap^3G-only revertants suggested that they restored at least some replication deficiencies of the ^cap^3G-only virus. In fact, gRNA packaging was significantly improved for all tested ^cap^3G-only revertants. We observed a gradient of packaging improvement in the revertants, such that packaging for the original ^cap^3G-only virus < ^cap^3G-R1 < ^cap^3G-R2 < ^cap^3G-R3, with gRNA packaging in ^cap^3G-R3 restored to WT levels. These differences may explain why the rapidly appearing ^cap^3G-R1 revertant was displaced by ^cap^3G-R2 and ^cap^3G-R3 at later time points.

Splicing, which was increased relative to WT in the ^cap^3G-only virus, was also restored to near-WT levels in the revertants, with the highest proportion of unspliced RNA observed in ^cap^3G-R3. All HIV-1 RNA splicing initiates with the use of the same 5′ splice site, termed D1, which is regulated at least in part by local secondary structure [27, 28]. The alternative secondary structures adopted by ^cap^1G and ^cap^3G RNA 5′ leaders are predicted to differ in D1 accessibility. Restored packaging and splicing levels, i.e. a shift of gRNA from the splicing/translation pool to the packaging pool, accompanied and may explain rapid spread of the revertants 3GR2 and 3GR3 in the infected culture. Moreover, when TSS sequences of ^cap^3G-R2 and ^cap^3G-R3 were cloned into an infectious virus background, the revertants were observed to replicate with kinetics similar to wild type.

Although none of the reversions restored the wild type TSS sequence, all identified ^cap^3G-only revertants acquired the use of multiple transcription start sites and displayed a packaging bias for one or a subset of their RNA isoforms. However, these selection preferences were not readily predictable. For example, the ^cap^GGA and ^cap^A RNAs produced by the ^cap^3G-R2 revertant correspond in size to ^cap^3G and ^cap^1G RNAs, but ^cap^GGA RNA and not ^cap^A RNA was enriched in virions. This suggests that the single 5′ A is deleterious to the packaging function of this RNA and possibly its folded form. Recent findings have underscored the importance of fine-tuning alternate 5′ leader structure stability to the functional roles of HIV-1 transcripts [22].

Heterogeneous TSS usage is a highly conserved innovation of the HIV-1 lineage [8, 9], and the studies here confirmed that the 5′ ends of HIV-1 RNAs dictate complementary RNA replication functions. The fact that ^cap^3G-only virus revertants evolved to acquire TSS use, splicing, and packaging properties similar to WT suggests that optimizing these processes is important to viral replication success. However, the benefits of expressing ^cap^3G RNA were less clear because ^cap^1G-only viruses replicated at rates similar to WT, and the only ^cap^1G-only revertant detected did not enhance replication much if at all. Nonetheless, the conservation of heterogeneous TSS use suggests that ^cap^3G RNA is beneficial to virus replication under conditions not captured by the experimental approaches here. One possibility would be during the initial expression of proviral DNA when enhancing the level of spliced RNAs might increase Tat and Rev expression and promote more robust expression. Thus, this work leaves unresolved a complete understanding of the selective advantages for the highly conserved function of expressing two isoforms of HIV-1 RNA.

## Materials and methods

### Plasmids, HIV-1 vectors and helpers

Previously published plasmids are as follows: the replication defective vector that included the HIV-1 NL4-3 strain RNA leader plus *gag, gag-pol, tat* and *rev* genes with puromycin resistance cassette has previously been referred to as HIV-1 GPP [5]; Minimal vector: NL4-3 based vector containing two LTRs, the 5′ leader, RRE and puromycin cassette; previously referred to as HIV-1 Native [5]; CMVβR8.2, a Ψ-HIV-1 helper [29]; and pNL4-3, infectious NL4-3 molecular clone [30]. ^cap^1G-only and ^cap^3G-only variants of Minimal were described previously [8] and used to template PCR fragments subcloned into HIV-1 GPP or pNL4-3 to generate ^cap^1G-only and ^cap^3G-only variants. Revertant sequences PCR amplified from cell DNA as described below were cloned into pCR4-TOPO (ThermoFisher) and subsequently cloned into Minimal vectors or pNL4-3. Minimal Δ variants contained a 94 b deletion upstream of the puromycin resistance gene and were created by near full-length plasmid amplification and subsequent self-ligation.

Full-length self-tagging viruses for single viral particle florescent microscopy assay were derived from a version of pNL4-3 [30] modified to carry inactivating mutations in *env*, *vpr*, and *nef* (E-R-Luc). The mVenus reading frame was inserted into *gag* between the sequences encoding for the Gag Matrix (MA) and Capsid (CA) domains [31]. Twenty-four copies of the MS2 bacteriophage stem loop [32] were inserted into the *pol* open reading frame downstream of the *gag* stop codon [33]. To detect the RNA through binding to the MS2 stem loops, cDNA encoding an MS2-mCherry fusion protein and harboring an SV40 nuclear localization signal was inserted into the *nef* open reading frame replacing the luciferase reporter, using NotI and XhoI restriction sites. HIV-1 promoter variants [8] were introduced into pNL4-3 Gag-mVenus/24xMSL/MS2-mCherry two-color self-tagging proviral plasmids using AatII and SpeI sites. All plasmids were verified using diagnostic restriction digestion and sequencing.

### Cells, viruses, transfection, virus release assays and infections

Human embryonic kidney 293T cells were purchased from the American Type Culture Collection (ATCC, Manassas, VA, USA). MT-4-EGFP cells were kindly provided by P. Bieniasz. To express HIV-1 vectors or produce infectious HIV-1 particles, freshly seeded 293T cells were grown in DMEM supplemented with 10% fetal bovine serum (FBS) and 50 µg/mL gentamicin at 37°C with 5% CO_2_ and transfected using polyethylenimine (Polysciences) [34]. Minimal vectors were co-transfected with CMVΔR8.2 at a 2:1 molar ratio, and HIV-GPP derivatives at a 1:1 molar ratio (8 µg of plasmid DNA total). For single cycle infectivity assays, HIV-GPP derivatives (4 µg) were co-transfected with 1 µg of vesicular stomatitis virus (VSV) G protein expression plasmid (pHEF-VSVG) [35]. For infectious HIV-1 derivatives, 5 µg of plasmid DNA was used for transfection.

Viral particle production was monitored by p24 enzyme-linked immunosorbent assay (ELISA) and/or quantitative PCR-based RT assay [36]. HIV-1 containing medium with a known concentration of CA-p24 was used as the standard. Viral infectivity was determined by puromycin resistant colony forming units/ml as described previously [37].

MT-4-eGFP cells were grown in RPMI supplemented with 10% FBS, 50 µg/mL gentamicin and 1.25 µg/mL puromycin at 37°C with 5% CO_2_ in 25cm^2^ culture flasks. To establish chronically infected MT-4-eGFP cells, viral media containing 2.5 ng of CA-p24 were added to 2 × 10^6^ freshly seeded MT-4-eGFP cells. Aliquots of the media were taken every 2nd or 3rd day of the infected cell passaging and analyzed by quantitative RT assay [36].

### Primary T Cell Isolation and Infection

Peripheral blood mononuclear cells (PBMCs) were isolated from fresh blood from anonymous healthy donors provided by the New York Blood Center using Ficoll-Paque PLUS (Cytiva) centrifugation and SepMate tubes (Stemcell Technologies) according to the manufacturer’s protocol. Total CD4+ T cells were isolated from PBMCs using a CD4+ T Cell Isolation Kit, human (Miltenyi Biotec) according to the manufacturer’s protocol. Isolated CD4+ T cells were maintained in RPMI supplemented with 10% FBS, 0.33 ug/mL amphotericin B, 50 ug/mL gentamicin, 1 mM sodium pyruvate, 1X GlutaMAX, 10 mM HEPES, and 1X NEAA (Gibco). The cells were stimulated using 6 ug/mL Phytohemagglutinin (PHA) (Thermo Scientific) in the presence of 10 ng/mL recombinant human IL-2 and 10 ng/mL of recombinant human IL-15 (BioLegend) for 3 days. On day 2 of activation, the cells were infected with virus in 0.4 ug/mL polybrene by spinoculation at 2500 rpm for 2 hours at room temperature in the 6-well culture plate (Corning Incorporated) with virus equivalent of 25ng of p24 per 3 x 10^6^ CD4+ cells per well. After spinoculation cells were washed twice with PBS and seeded into 25cm^2^ culture flasks.

### Microscopy and image analysis

To generate labeled virus-like particles, approximately 500,000 HEK293T cells were plated in each well of a 6-well dish and transfected with plasmids encoding the wild-type, 1G-only, or 3G-only two-color self-tagging viruses using polyethylenimine (PEI). The media was exchanged at 24-hours post-transfection and virus particle-containing supernatants were harvested at 48-hours post-transfection, filtered through a 0.45μm filter, and centrifuged through 20% sucrose for 2 hours at 15,000 rpm. The medium was discarded after centrifugation and concentrated viral particles were resuspended in 1xPBS, plated in a 24-well glass-bottom dish (Cellvis, Mountain View, CA), and left overnight at 4°C to allow virus particles to settle down on the glass wells. Microscopy was performed using a Nikon Ti-Eclipse inverted wide-field microscope (Nikon Corp, Minato, Tokyo, Japan) using a 100x Plan Apo oil objective lens (numerical aperture [NA] 1.45). Cell and virion images were captured using an ORCA-Flash4.0 CMOS camera (Hamamatsu Photonics, Skokie, IL, USA) and the following excitation/emission filter sets: 510/535nm (YFP) and 585/610nm (mCherry). All images were processed and analyzed using FIJI/ImageJ2 [38].

The Cellpose TrackMate plugin [39] was used to measure mean fluorescent intensities (MFIs). Spot IDs were created based on MS2-mCherry signal masks and applied to the Gag-YFP channel to generate a per cell fluorescent profile. A custom FIJI/ImageJ2 workflow (https://github.com/elevans/dbp-solutions/blob/main/scripts/sherer/sl_sva.py) was used to threshold particles and create masks that encompassed the virion YFP fluorescence, corresponding to the per virion signal from Gag-YFP structural protein. Using those masks, the mCherry fluorescence corresponding to the US RNA (MS2-mCherry) signal was measured. All signals (cell and virion) were background-subtracted using a negative transfection control prior to quantitative analyses. Cell and virion background subtracted MFIs for MS2-mCherry and Gag-YFP channels were plotted using GraphPad Prism (version 10.3.1). Outliers were identified and removed using the ROUT method and Q=1% aggression. The cleaned data was plotted for MS2-mCherry MFI, Gag-YFP MFI, and, for virions, MS2-mCherry/Gag-YFP MFI. A one-way analysis of variance (ANOVA) using multiple comparisons was performed to determine statistically significant differences between the means of MS2-mCherry MFIs and Gag-YFP MFIs, and MS2-mCherry/Gag-YFP ratios.

### RNA and DNA extraction and analysis

Viral particles were pelleted by ultracentrifugation of filtered viral media at 25000 RPM for 2h. RNA was extracted from pelleted virions and cells with TRIzol (Invitrogen) according to manufacturer protocol. RNA samples were treated with RQ1 DNase (Promega) and re-extracted with phenol/chloroform. RNase Protection Assays (RPA) were performed as described [40]. Riboprobes used in this study: HIV*gag*/7SL, a chimeric riboprobe targeting HIV-1 *gag* (200bp) and host 7SL RNA (100bp), HIV*gag*/CMV, targeting *gag* (200bp) in NL4-3 GPP derivatives and CMV promoter region in the Minimal vectors (289bp in the Minimal vector and 195bp in the Minimal Δ vector); HIV unspliced/spliced, riboprobe targeting D1 region in the HIV-1 leader, protecting 130 bp in unspliced and 60 bp in spliced HIV-1 RNA. Dried RPA gels were quantified by phosphorimaging with ImageQuant TL 10.2 software.

HIV-1 derivatives’ RNAs’ 5’ ends were analyzed by CaDAL assay [8] using the TeloPrime Full-Length cDNA Amplification Kit V2 (Lexogen) components as described previously [8].

For proviral DNA analysis, cellular genomic DNA was isolated using the DNeasy Tissue and Blood kit (Qiagen, Valencia, CA) according to the manufacturer protocol. A portion of proviral DNA including most of the 5’LTR and the entire *gag* gene (approximately 2.2 kb) was amplified with primers GCTAATTCACTCCCAAAGAAGACAAG (forward) and CAAACCTGAAGCTCTCTTCTGGTG (reverse) and Phusion polymerase (NEB). PCR products were extracted from agarose using the NEB Monarch DNA gel extraction kit and used for Sanger sequencing. For individual molecular clones, PCR products were cloned using a TOPO TA kit (Thermo Fisher) and then sequenced.

### High-throughput sequencing and data analysis

RNA samples were extracted from the cells 3 days post-infection. As a no-replication control sample, RNA was extracted from cells infected with WT NL4-3 virus in the presence of the two antiretroviral drugs (ARD) AZT and reltagravir. 2.2 kb PCR products obtained by amplification of proviral DNA from infected cells (see RNA and DNA extraction and analysis section) was used as a template for a secondary PCR with following primers: ACACTCTTTCCCTACACGACGCTCTTCCGATCTGACATCGAGCTTGCTACAAGGGAC (forward, specific to HIV-1 U3, 125 bp upstream of TSS) and GACTGGAGTTCAGACGTGTGCTCTTCCGATCTGAGGGATCTCTAGTTACCAGAGTCAC (reverse, specific to HIV-1 U5 sequence 147bp downstream of TSS). Besides HIV-1 specific sequences, both primers included Illumina partial adapters sequences. PCR products were sent to GENEWIZ (South Plainsfield, NJ, USA) for sequencing (Amplicon EZ service) using an Illumina MiSeq platform and 250-bp paired-end reads. TSS reversions were analyzed using an in-house script, available upon request from the Telesnitsky lab.

Deep sequencing splicing analysis was done using a protocol from Emery *et al* 2017 [21] with the following adaptations. In separate reactions, two cDNA primers were used. GTGCTCTTCCGATCTNNNNNNNNNNNNNN has 14 random bases that serve as a Unique Molecular Identifier (UMI) as well as a universal primer. GTGCTCTTCCGATCTNNNNNNNNNNTTTYCCACCCCC has a 10-base random UMI and a sequence that primes at two regions of the HIV NL4-3 genome, 6257 and 8576, downstream of splice sites D4 and A7 respectively. All of the bead purified cDNA product was used as input to the first PCR step. The semi-nested first PCR step used a forward primer upstream of D1 (ATCTCTCGACGCAGGAC) and this reverse primer (TTCAGACGTGTGCTCTTCCGATCT). 5 μl of this bead purified first PCR was used as input to a second PCR, which used forward primer (GCCTCCCTCGCGCCATCAGAGATGTGTATAAGAGACAGNNNNTGCTGAAGCGCGCACGGCA AG) and reverse primer (TTCAGACGTGTGCTCTTCCGATCT). 5 μl of this bead purified second PCR was used as input to the final PCR, which used the primers previously described [21] to add Illumina platform sequences. Thermocycler settings for all PCR reactions were: 95°C initial denaturing for 5 min; then 3x cycles with 95°C 30 sec, annealing for 15 sec at 72°C, extension for 2 min at 72°C; then 3x each with decreasing annealing temps at 70, 68, 66, 64, and 62°C; ending with 12 (nested) or 17 (final PCR) cycles with annealing temp at 60°C. A detailed and user-friendly protocol is available from the Swanstrom lab. Sequencing was done using Illumina MiSeq 300 paired end reads and this Illumina primer: GCCTCCCTCGCGCCATCAGAGATGTGTATAAGAGACAG. The Illumina bcl2fastq pipeline was used (v.2.20.0) for initial processing of data. Splice site quantification was done using in house programs available from the Swanstrom lab.

### HIV-1 protein analysis

293T cells were lysed in RIPA buffer [150 mM NaCl, 50 mM Tris pH 7.5, 1% NP40, 0.5% Deoxycholate, 0.1% SDS], samples were separated via SDS-PAGE and transferred to Immun-Blot PVDF Membrane (Bio-Rad), blocked in 1% milk in 1x TBS and incubated with Human (HIV-IG) (NIH-ARP, 3957) and anti-β-actin mouse (Invitrogen, AM4302) in 1X TBST. After washing, the membrane was incubated with secondary antibodies: goat anti-mouse IRDye 680RD (LI-COR, 926-68070) and goat anti-human IRDye 800cw (LI-COR, 925-32232). Finally, the immunoblot was imaged using an Amersham Typhoon (Cytiva). Gag/β-actin ratios were quantified using ImageQuant TL 10.2 software.

## Acknowledgements

We acknowledge Jordan Becker for engineering the prototype self-tagging genome, Edward Evans III for assisting computational analysis for SVA, and Emily Schugardt for critically reading the manuscript. We also acknowledge support from R01 AI50498 to AT, R56 AI110221 to NS, and from U54 AI170660 to AT, RS and NS. KG and SC received support for this work from training grants T32 GM145470 and T32 AI007528, respectively.

## References

1. Kharytonchyk, S., Monti, S., Smaldino, P. J., Van, V., Bolden, N. C., Brown, J. D., Russo, E., Swanson, C., Shuey, A., Telesnitsky, A. & Summers, M. F. (2016). Transcriptional start site heterogeneity modulates the structure and function of the HIV-1 genome. Proc Natl Acad Sci U S A. 113, 13378–13383. 10.1073/pnas.1616627113.

2. Masuda, T., Sato, Y., Huang, Y. L., Koi, S., Takahata, T., Hasegawa, A., Kawai, G. & Kannagi, M. (2015). Fate of HIV-1 cDNA intermediates during reverse transcription is dictated by transcription initiation site of virus genomic RNA. Sci Rep. 5, 17680. 10.1038/srep17680.

3. Esquiaqui, J. M., Kharytonchyk, S., Drucker, D. & Telesnitsky, A. (2020). HIV-1 spliced RNAs display transcription start site bias. RNA. 26, 708–714. 10.1261/rna.073650.119.

4. Brown, J. D., Kharytonchyk, S., Chaudry, I., Iyer, A. S., Carter, H., Becker, G., Desai, Y., Glang, L., Choi, S. H., Singh, K., Lopresti, M. W., Orellana, M., Rodriguez, T., Oboh, U., Hijji, J., Ghinger, F. G., Stewart, K., Francis, D., Edwards, B., Chen, P., Case, D. A., Telesnitsky, A. & Summers, M. F. (2020). Structural basis for transcriptional start site control of HIV-1 RNA fate. Science. 368, 413–417. 10.1126/science.aaz7959.

5. Ding, P., Kharytonchyk, S., Kuo, N., Cannistraci, E., Flores, H., Chaudhary, R., Sarkar, M., Dong, X., Telesnitsky, A. & Summers, M. F. (2021). 5’-Cap sequestration is an essential determinant of HIV-1 genome packaging. Proc Natl Acad Sci U S A. 118. 10.1073/pnas.2112475118.

6. Chen, J., Nikolaitchik, O., Singh, J., Wright, A., Bencsics, C. E., Coffin, J. M., Ni, N., Lockett, S., Pathak, V. K. & Hu, W. S. (2009). High efficiency of HIV-1 genomic RNA packaging and heterozygote formation revealed by single virion analysis. Proc Natl Acad Sci U S A. 106, 13535–13540. 10.1073/pnas.0906822106.

7. Nikolaitchik, O. A., Dilley, K. A., Fu, W., Gorelick, R. J., Tai, S. H., Soheilian, F., Ptak, R. G., Nagashima, K., Pathak, V. K. & Hu, W. S. (2013). Dimeric RNA recognition regulates HIV-1 genome packaging. PLoS Pathog. 9, e1003249. 10.1371/journal.ppat.1003249.

8. Kharytonchyk, S., Burnett, C., Gc, K. & Telesnitsky, A. (2023). Transcription start site heterogeneity and its role in RNA fate determination distinguish HIV-1 from other retroviruses and are mediated by core promoter elements. J Virol. 97, e0081823. 10.1128/jvi.00818-23.

9. Nikolaitchik, O. A., Islam, S., Kitzrow, J. P., Duchon, A., Cheng, Z., Liu, Y., Rawson, J. M. O., Shao, W., Nikolaitchik, M., Kearney, M. F., Maldarelli, F., Musier-Forsyth, K., Pathak, V. K. & Hu, W. S. (2023). HIV-1 usurps transcription start site heterogeneity of host RNA polymerase II to maximize replication fitness. Proc Natl Acad Sci U S A. 120, e2305103120. 10.1073/pnas.2305103120.

10. Yoshida, T., Kasuya, Y., Yamamoto, H., Kawai, G., Hanaki, K. i., Matano, T. & Masuda, T. (2024). HIV-1 RNAs whose transcription initiates from the third deoxyguanosine of GGG tract in the 5’ long terminal repeat serve as a dominant genome for efficient provirus DNA formation. J Virol. 98, e0182523. 10.1128/jvi.01825-23.

11. Rawson, J. M. O., Nikolaitchik, O. A., Shakya, S., Keele, B. F., Pathak, V. K. & Hu, W. S. (2022). Transcription Start Site Heterogeneity and Preferential Packaging of Specific Full-Length RNA Species Are Conserved Features of Primate Lentiviruses. Microbiol Spectr. 10, e0105322. 10.1128/spectrum.01053-22.

12. Clavel, F. & Orenstein, J. M. (1990). A mutant of human immunodeficiency virus with reduced RNA packaging and abnormal particle morphology. J Virol. 64, 5230–5234. 10.1128/JVI.64.10.5230-5234.1990.

13. Heng, X., Kharytonchyk, S., Garcia, E. L., Lu, K., Divakaruni, S. S., LaCotti, C., Edme, K., Telesnitsky, A. & Summers, M. F. (2012). Identification of a minimal region of the HIV-1 5’-leader required for RNA dimerization, NC binding, and packaging. J Mol Biol. 417, 224–239. 10.1016/j.jmb.2012.01.033.

14. Kharytonchyk, S., Brown, J. D., Stilger, K., Yasin, S., Iyer, A. S., Collins, J., Summers, M. F. & Telesnitsky, A. (2018). Influence of gag and RRE Sequences on HIV-1 RNA Packaging Signal Structure and Function. J Mol Biol. 430, 2066–2079. 10.1016/j.jmb.2018.05.029.

15. Rulli, S. J., Jr., Hibbert, C. S., Mirro, J., Pederson, T., Biswal, S. & Rein, A. (2007). Selective and nonselective packaging of cellular RNAs in retrovirus particles. J Virol. 81, 6623–6631. 10.1128/JVI.02833-06.

16. Laham-Karam, N. & Bacharach, E. (2007). Transduction of human immunodeficiency virus type 1 vectors lacking encapsidation and dimerization signals. J Virol. 81, 10687–10698. 10.1128/JVI.00653-07.

17. Houzet, L., Paillart, J. C., Smagulova, F., Maurel, S., Morichaud, Z., Marquet, R. & Mougel, M. (2007). HIV controls the selective packaging of genomic, spliced viral and cellular RNAs into virions through different mechanisms. Nucleic Acids Res. 35, 2695–2704. 10.1093/nar/gkm153.

18. Das, A. T., Vrolijk, M. M., Harwig, A. & Berkhout, B. (2012). Opening of the TAR hairpin in the HIV-1 genome causes aberrant RNA dimerization and packaging. Retrovirology. 9, 59. 10.1186/1742-4690-9-59.

19. Fernandez, M. V., Hoffman, H. K., Pezeshkian, N., Tedbury, P. R., van Engelenburg, S. B. & Freed, E. O. (2020). Elucidating the Basis for Permissivity of the MT-4 T-Cell Line to Replication of an HIV-1 Mutant Lacking the gp41 Cytoplasmic Tail. J Virol. 94. 10.1128/JVI.01334-20.

20. Harada, S., Koyanagi, Y. & Yamamoto, N. (1985). Infection of HTLV-III/LAV in HTLV-I-carrying cells MT-2 and MT-4 and application in a plaque assay. Science. 229, 563–566. 10.1126/science.2992081.

21. Emery, A., Zhou, S., Pollom, E. & Swanstrom, R. (2017). Characterizing HIV-1 Splicing by Using Next-Generation Sequencing. J Virol. 91. 10.1128/JVI.02515-16.

22. Yasin, S., Lesko, S. L., Kharytonchyk, S., Brown, J. D., Chaudry, I., Geleta, S. A., Tadzong, N. F., Zheng, M. Y., Patel, H. B., Kengni, G., Neubert, E., Quiambao, J. M. C., Becker, G., Ghinger, F. G., Thapa, S., Williams, A., Radov, M. H., Boehlert, K. X., Hollmann, N. M., Singh, K., Bruce, J. W., Marchant, J., Telesnitsky, A., Sherer, N. M. & Summers, M. F. (2024). Role of RNA structural plasticity in modulating HIV-1 genome packaging and translation. Proc Natl Acad Sci U S A. 121, e2407400121. 10.1073/pnas.2407400121.

23. Cheng, Z., Islam, S., Kanlong, J. G., Sheppard, M., Seo, H., Nikolaitchik, O. A., Kearse, M. G., Pathak, V. K., Musier-Forsyth, K. & Hu, W. S. (2024). Translation of HIV-1 unspliced RNA is regulated by 5’ untranslated region structure. J Virol. e0116024. 10.1128/jvi.01160-24.

24. Kulpa, D., Topping, R. & Telesnitsky, A. (1997). Determination of the site of first strand transfer during Moloney murine leukemia virus reverse transcription and identification of strand transfer-associated reverse transcriptase errors. EMBO J. 16, 856–865. 10.1093/emboj/16.4.856.

25. Swanstrom, R., Varmus, H. E. & Bishop, J. M. (1981). The terminal redundancy of the retrovirus genome facilitates chain elongation by reverse transcriptase. J Biol Chem. 256, 1115–1121.

26. Volloch, V. Z., Schweitzer, B. & Rits, S. (1995). Transcription of the 5’-terminal cap nucleotide by RNA-dependent DNA polymerase: possible involvement in retroviral reverse transcription. DNA Cell Biol. 14, 991–996. 10.1089/dna.1995.14.991.

27. Mueller, N., Das, A. T. & Berkhout, B. (2016). A Phylogenetic Survey on the Structure of the HIV-1 Leader RNA Domain That Encodes the Splice Donor Signal. Viruses. 8. 10.3390/v8070200.

28. Mueller, N., Klaver, B., Berkhout, B. & Das, A. T. (2015). Human immunodeficiency virus type 1 splicing at the major splice donor site is controlled by highly conserved RNA sequence and structural elements. J Gen Virol. 96, 3389–3395. 10.1099/jgv.0.000288.

29. Naldini, L., Blomer, U., Gallay, P., Ory, D., Mulligan, R., Gage, F. H., Verma, I. M. & Trono, D. (1996). In vivo gene delivery and stable transduction of nondividing cells by a lentiviral vector. Science. 272, 263–267. 10.1126/science.272.5259.263.

30. Adachi, A., Gendelman, H. E., Koenig, S., Folks, T., Willey, R., Rabson, A. & Martin, M. A. (1986). Production of acquired immunodeficiency syndrome-associated retrovirus in human and nonhuman cells transfected with an infectious molecular clone. J Virol. 59, 284–291. 10.1128/JVI.59.2.284-291.1986.

31. Behrens, R. T., Rajashekar, J. K., Bruce, J. W., Evans, E. L., 3rd, Hansen, A. M., Salazar-Quiroz, N., Simons, L. M., Ahlquist, P., Hultquist, J. F., Kumar, P. & Sherer, N. M. (2023). Exploiting a rodent cell block for intrinsic resistance to HIV-1 gene expression in human T cells. mBio. 14, e0042023. 10.1128/mbio.00420-23.

32. Femino, A. M., Fay, F. S., Fogarty, K. & Singer, R. H. (1998). Visualization of single RNA transcripts in situ. Science. 280, 585–590. 10.1126/science.280.5363.585.

33. Pocock, G. M., Becker, J. T., Swanson, C. M., Ahlquist, P. & Sherer, N. M. (2016). HIV-1 and M-PMV RNA Nuclear Export Elements Program Viral Genomes for Distinct Cytoplasmic Trafficking Behaviors. PLoS Pathog. 12, e1005565. 10.1371/journal.ppat.1005565.

34. Boussif, O., Lezoualc’h, F., Zanta, M. A., Mergny, M. D., Scherman, D., Demeneix, B. & Behr, J. P. (1995). A versatile vector for gene and oligonucleotide transfer into cells in culture and in vivo: polyethylenimine. Proc Natl Acad Sci U S A. 92, 7297–7301. 10.1073/pnas.92.16.7297.

35. Chang, L. J., Urlacher, V., Iwakuma, T., Cui, Y. & Zucali, J. (1999). Efficacy and safety analyses of a recombinant human immunodeficiency virus type 1 derived vector system. Gene Ther. 6, 715–728. 10.1038/sj.gt.3300895.

36. Vermeire, J., Naessens, E., Vanderstraeten, H., Landi, A., Iannucci, V., Van Nuffel, A., Taghon, T., Pizzato, M. & Verhasselt, B. (2012). Quantification of reverse transcriptase activity by real-time PCR as a fast and accurate method for titration of HIV, lenti- and retroviral vectors. PLoS One. 7, e50859. 10.1371/journal.pone.0050859.

37. Kharytonchyk, S., King, S. R., Ndongmo, C. B., Stilger, K. L., An, W. & Telesnitsky, A. (2016). Resolution of Specific Nucleotide Mismatches by Wild-Type and AZT-Resistant Reverse Transcriptases during HIV-1 Replication. J Mol Biol. 428, 2275–2288. 10.1016/j.jmb.2016.04.005.

38. Schindelin, J., Arganda-Carreras, I., Frise, E., Kaynig, V., Longair, M., Pietzsch, T., Preibisch, S., Rueden, C., Saalfeld, S., Schmid, B., Tinevez, J. Y., White, D. J., Hartenstein, V., Eliceiri, K., Tomancak, P. & Cardona, A. (2012). Fiji: an open-source platform for biological-image analysis. Nat Methods. 9, 676–682. 10.1038/nmeth.2019.

39. Stringer, C., Wang, T., Michaelos, M. & Pachitariu, M. (2021). Cellpose: a generalist algorithm for cellular segmentation. Nat Methods. 18, 100–106. 10.1038/s41592-020-01018-x.

40. Onafuwa-Nuga, A. A., King, S. R. & Telesnitsky, A. (2005). Nonrandom packaging of host RNAs in moloney murine leukemia virus. J Virol. 79, 13528–13537. 10.1128/JVI.79.21.13528-13537.2005.

